# Selective Labeling and Identification of the Tumor Cell Proteome of Pancreatic Cancer *In Vivo*

**DOI:** 10.1101/2020.05.25.113670

**Authors:** Nancy G. Azizian, Delaney K. Sullivan, Litong Nie, Sammy Pardo, Dana Molleur, Junjie Chen, Susan T. Weintraub, Yulin Li

## Abstract

Pancreatic ductal adenocarcinoma (PDAC) is among the deadliest cancers. Dissecting the tumor cell proteome from that of the non-tumor cells in the PDAC tumor bulk is critical for tumorigenesis studies, biomarker discovery, and development of therapeutics. However, investigating the tumor cell proteome has proven evasive due to the tumor’s extremely complex cellular composition. To circumvent this technical barrier, we have combined bioorthogonal non-canonical amino acid tagging (BONCAT) and data-independent acquisition mass spectrometry (DIA-MS) in an orthotopic PDAC model to specifically identify the tumor cell proteome *in vivo*. Utilizing the tumor cell-specific expression of a mutant tRNA synthetase transgene, this approach provides tumor cells with the exclusive ability to incorporate an azide-bearing methionine analog into newly synthesized proteins. The azide-tagged tumor cell proteome is subsequently enriched and purified via a bioorthogonal reaction and then identified and quantified using DIA-MS. Applying this workflow to the orthotopic PDAC model, we have identified thousands of proteins expressed by the tumor cells. Furthermore, by comparing the tumor cell and tumor bulk proteomes, we showed that the approach can distinctly differentiate proteins produced by tumor-cells from non-tumor cells within the tumor microenvironment. Our study, for the first time, reveals the tumor cell proteome of PDAC under physiological conditions, providing broad applications for tumorigenesis, therapeutics, and biomarker studies in various human cancers.

## Introduction

The tumor bulk is not only a mass of proliferating tumor cells, but also consists of a variety of non-tumor cells, secreted factors, and the extracellular matrix, which are collectively known as the tumor microenvironment (TME). The interaction between tumor cells and the surrounding TME has profound impact on all stages of tumor development. Human pancreatic ductal adenocarcinoma (PDAC), in particular, has a highly complex TME, imparted by a dense desmoplastic stroma and a host of stromal fibroblasts, endothelial, inflammatory, and immune cells. The stromal components of human PDAC may account for up to 80% of the total tumor volume, with tumor cells constituting a minor population.^1–3^ Dissecting proteins produced by the PDAC tumor cells from those of non-tumor cells in the TME is critical for tumorigenesis and therapeutic studies. However, the heterogeneous and complex cellular composition of the PDAC tumor mass, has thus far, precluded precise isolation and identification of the PDAC tumor cell proteome *in vivo*.

Direct investigation of the PDAC tumor cell proteome requires selective purification of proteins from the tumor cells and not the non-tumor cells within the tumor bulk. Several recent studies have focused on cell-selective metabolic labeling of the proteomes.^4^ These approaches include cell-type-specific labeling using amino acid precursors (CTAP),^5^ bioorthogonal non-canonical amino acid tagging (BONCAT),^6^ and stochastic orthogonal recording of translation (SORT).^7^ BONCAT has been shown to label cell-selective proteomes in the fruit fly,^8^ as well as mouse brain and muscle.^9–11^ BONCAT works through bioorthogonal chemical reactions that do not exist in nature, and, thus, will not cross-react with any physiological processes in the cells.^12,13^ This technique relies on bioorthogonal incorporation of azide-bearing methionine analogs, such as azidonorleucine (ANL) and azidohomoalanine (AHA), into newly synthesized polypeptides. Due to the small size of the azide moiety, ANL or AHA incorporation has no apparent effect on protein function.^9–11^ During protein translation, ANL is preferentially recognized and charged onto tRNA^Met^ by a mutant methionyl-tRNA synthetase (MetRS^L274G^), and is subsequently incorporated in the elongating polypeptide chains (**Fig 1A-B**).^14^ ANL-tagged proteins can be selectively conjugated and enriched through azide-alkyne cycloaddition.^13^ Further identification of the proteome is achieved through mass spectrometric (MS) analysis of the ANL-tagged proteins (**Fig 1C**). ANL incorporation is unbiased, non-toxic, biocompatible, and does not affect protein stability.^6^

**Figure 1.**
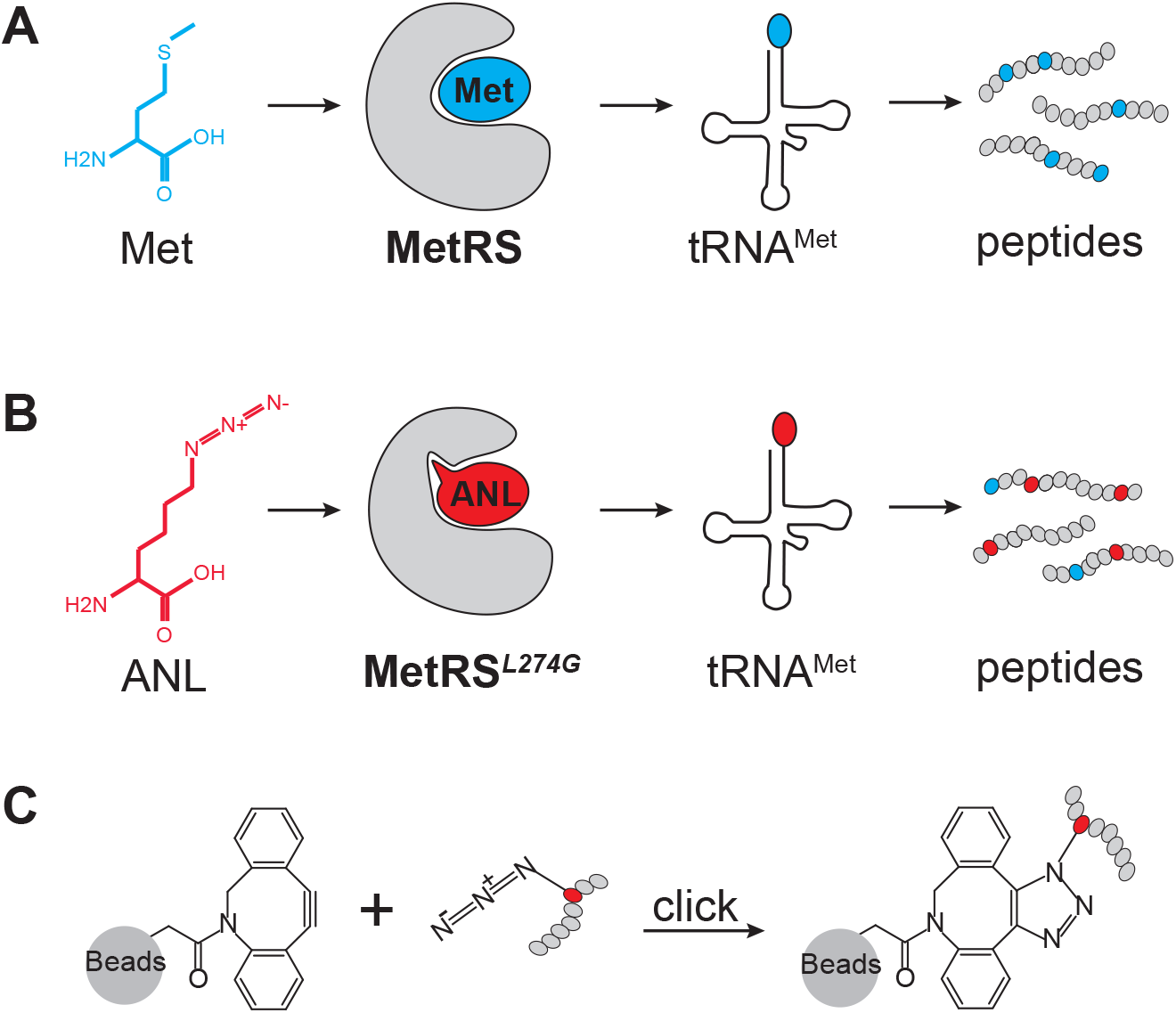
MetRS^L274G^-assisted peptide labeling and BONCAT reaction. (A-B) Diagram showing cell selective labeling of proteome by BONCAT. (C) Diagram showing selective coupling of ANL labeled peptides using DBCO-alkyne beads.

The PDAC bulk tumor is composed of tumor cells and many types of non-tumor cells. The ectopic expression of MetRS^L274G^ transgene in tumor cells, but not the non-tumor cells in the tumor bulk, enables the exclusive tagging of the tumor cell proteome using ANL. Notably, the absence of the MetRS^L274G^ transgene in the various non-tumor cells in the TME precludes ANL incorporation into their proteomes. Following ANL labeling, the tumor cell proteome is enriched and purified for MS analysis. Thus, applying BONCAT to the animal model of PDAC facilitates the identification of tumor cell proteome in a physiological context.

Data-independent acquisition mass spectrometry (DIA-MS) is a highly reproducible, state-of-the-art approach for quantitative proteomic analysis.^15–19^ Traditionally, data-dependent acquisition mass spectrometry (DDA-MS) has been used in a variety of label-free and label-based methods to measure quantitative changes in global protein levels in biological samples. However, the stochastic nature of DDA bears a bias toward higher abundance peptides.

Undersampling of medium and low abundance peptides causes inconsistencies in detection of peptides and hampers reproducibility among replicates. In the DIA-MS approach, all precursors are fragmented to yield tandem-MS data, providing sequence information from virtually all peptides in a sample with minimal loss of information. Due to its high accuracy and reproducibility, DIA-MS is a powerful method for comprehensive proteomic studies of complex samples, including tumor specimens.^20–22^

Here we have combined BONCAT bioorthogonal chemistry and DIA-MS proteomics to specifically investigate the tumor cell proteome in an orthotopic transplantation model of PDAC. We have identified over 3,000 proteins expressed in PDAC tumor cells, many of which are predominantly, if not exclusively, expressed in the tumor cells. Thus, we have established a robust technical platform for *in vivo* identification of the proteome of the tumor cells embedded within the bulk tumor, with broad applications in the studies of tumorigenesis, cancer therapeutics, and cancer detection.

## Results

### Construction and validation of PDAC-BONCAT cells

To label the proteome of PDAC tumor cells, we utilized the mutant murine methionyl-tRNA synthetase, MetRS^L274G^, which preferentially charges noncanonical amino acid azidonorleucine (ANL) to the elongator tRNA^Met^. ANL is utilized by MetRS^L274G^ and not by the wild-type translational machinery.^23^ In cells expressing MetRS^L274G^, polypeptide incorporation of ANL containing the reactive azide moiety enables selective conjugation to dyes and functionalized beads for visualization and enrichment.

The MetRS^L274G^ mutant transgene was cloned into a lentiviral vector and delivered to a murine pancreatic cancer cell line (4292)^24^ via lentiviral infection. Single-cell clones were derived and the expression of FLAG-tagged MetRS^L274G^ was confirmed by Western blot analysis (**Fig 2A**, **Supplementary Fig 1**). ANL incorporation by MetRS^L274G^ into the tumor cell proteome was visualized using the azide-reactive red-fluorescent tetramethyl rhodamine dibenzocyclooctyne (TAMRA-DBCO) alkyne probe. First, metabolic labeling was achieved by growing the tumor cells expressing MetRS^L274G^ in media containing ANL or control media for five hours. Cell lysates were separated by sodium dodecyl sulfate polyacrylamide gel electrophoresis (SDS-PAGE). Next, to perform in-gel fluorescence, TAMRA was covalently reacted onto the ANL azide moiety of the newly synthesized proteins via a copper-free click reaction. ANL-labeled proteins were detected through direct in-gel fluorescence (**Fig 2B**). The same SDS-PAGE gel was stained with Coomassie blue to visualize the total protein load and size distribution. Strong fluorescence signals were consistently detected in cells labeled with ANL, but not methionine (Met) (**Fig 2B-C**). Notably, ANL incorporation was evenly distributed across the proteome as judged by the similarity of the band patterns between TAMRA and Coomassie blue staining of the ANL-labeled samples (**Fig 2B-C**). Thus, with the cell-specific expression of the MetRS^L274G^ transgene, the PDAC-BONCAT system allows for effective and unbiased incorporation of ANL into the tumor cell proteome, facilitating subsequent enrichment and identification via mass-spectrometry.

**Figure 2.**
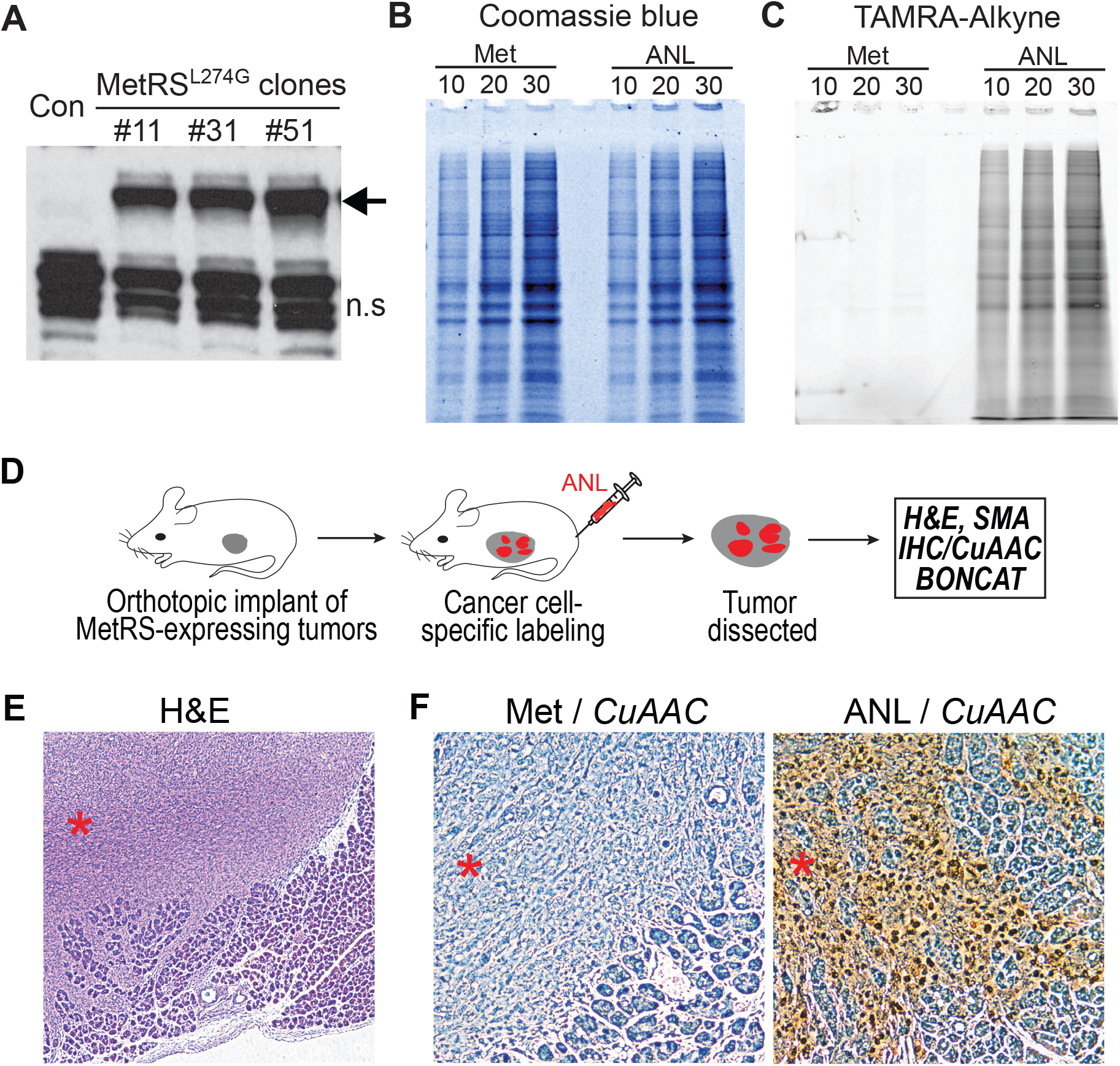
*in vitro* and *in vivo* validation of the PDAC-BONCAT system. (A) Western blot analysis of single-cell clones ectopically expressing the MetRS^L274G^ transgene. The MetRS^L274G^ protein is indicated by an arrow; n.s. indicates non-specific bands recognized by the FLAG antibody. (B-C) Detection of ANL-labeled proteins by TAMRA-DBCO and SDS-PAGE *in vitro*. 10, 20, and 30, indicate micrograms of protein lysate loaded per lane. (D) Overview of *in vivo* ANL labeling. (E) H&E staining showing tumor infiltration in the adjacent normal acinar tissues. Tumor nodule is indicated by a red asterisk. (F) IHC analysis of ANL incorporation by CuAAC. Met samples serve as the negative control. Tumor nodules are indicated by red asterisks.

### In vivo validation of PDAC-BONCAT system

To examine whether the PDAC-BONCAT system allows for *in vivo* tumor cell-specific proteome labeling, we set up an orthotropic transplantation model. The 4292 murine PDAC cells expressing the MetRS^L274G^ transgene were surgically implanted in the pancreata of immunodeficient *NOD-scid IL2R*γ^*null*^ (NSG) mice. These tumor-bearing mice were then randomized into ANL and Met groups. Mice in the ANL group were metabolically labeled via daily intraperitoneal (IP) injection of ANL for 10 days, while those in the Met group were injected with normal saline. Throughout the experiment, animals were provided with regular diet with no methionine depletion. At the end of the 10-day injection regimen, tumor samples were collected and processed for hematoxylin and eosin (H&E), and α-smooth muscle actin (α-SMA) immunohistochemistry (IHC) staining (**Fig 2D**). Additionally, *in situ* detection of ANL-incorporated proteins was performed using copper-catalyzed azide-alkyne cycloaddition (CuAAC).

H&E staining of the tumor sections revealed a highly heterogeneous population of cancer cells invading the adjacent acinar tissues (**Fig 2E**). IHC analysis with α-SMA antibody identified abundant stromal fibroblasts in the tumor bulk (**Supplementary Fig 2**). Collectively, these features confirm the establishment of a murine PDAC model, capable of recapitulating the heterogeneous cellular composition and histological features of human pancreatic cancer. Further *in situ* detection of ANL incorporation via CuAAC click reaction showed signals specifically localized in tumor cells but not in the adjacent normal cells and tissues (**Fig 2F**), confirming that the tumor cells but not the non-tumor cells in the tumor bulk can incorporate ANL into their proteome. Contrary to the ANL-labeled tissues, no signal was detected in tumor tissues isolated from animals in the Met group where ANL labeling had not taken place (**Fig 2F**). These *in vivo* observations demonstrate that BONCAT effectively tags the PDAC tumor cell proteome within its physiological milieu, and that the proteome labeling is highly specific to the tumor cells, distinguishing them from the various non-tumor cells in the TME.

### Defining the in vivo tumor proteome through coupling BONCAT and DIA-MS

For *in vivo* identification of tumor cell-specific proteins, surgical implantation of the PDAC-MetRS^L274G^ cells was performed in a large cohort of NSG mice. Following the same 10-day injection regimen for metabolic labeling, tumors from both ANL and Met groups were collected, lysed, and subjected to BONCAT purification and DIA-MS proteomic analysis.

TAMRA-alkyne cycloaddition reaction detected ANL incorporation in the tumor bulk lysates collected from the ANL but not the Met group (**Fig 3A-B**). Of note, the tumor bulk proteomes, represented by the relative intensities and overall pattern of the bands in lane 5-11 of Coomassie blue staining showed no correlation with the corresponding tumor-cell proteomes visualized by TAMRA staining in the same lanes. This observation suggests that the tumor bulk proteome, typically identified in preclinical and/or clinical analysis of tumor samples, does not adequately reflect the tumor cell proteome.

**Figure 3.**
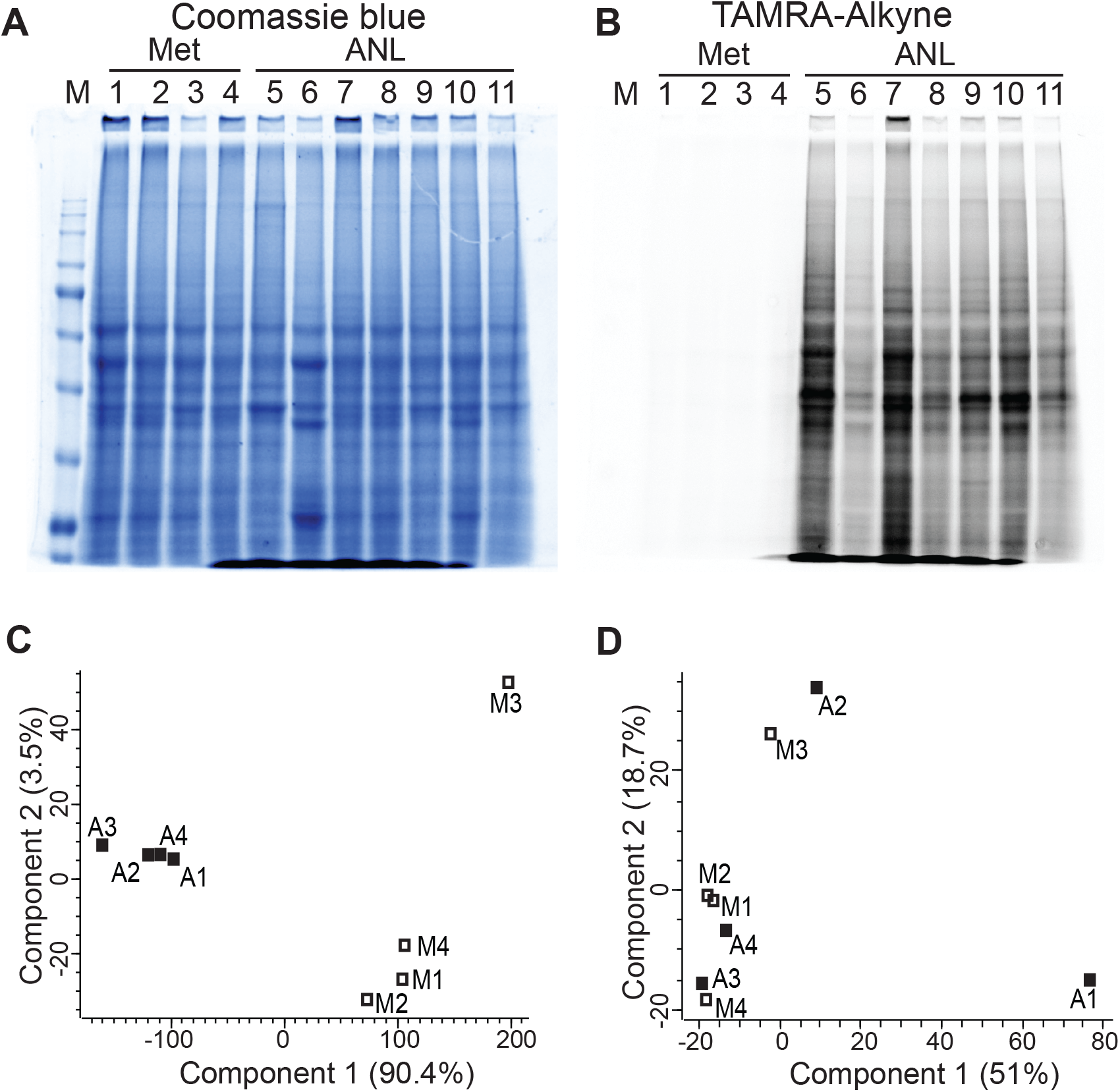
Quality analysis of BONCAT-enriched samples analyzed by DIA-MS. (A-B) TAMRA-alkyne and SDS-PAGE analysis of eleven *in vivo* tumor samples labeled with Met (lanes 1-4) or ANL (lanes 5-11). (C) PCA of DIA-MS data for BONCAT-ANL and BONCAT-Met samples. Four tumors are included in both the Met group (M1, 31C; M2, 30C; M3, 32C; and M4, 33C) and the ANL groups (A1, 40A; A2, ANL1; A3, 38A; and A4, 29A). (D) PCA of DIA-MS data for Bulk-ANL and Bulk-Met samples.

Four tumors from each group were randomly chosen for BONCAT enrichment and downstream DIA-MS proteomic analysis. Purification and enrichment of ANL-incorporated proteins in tumor lysates were achieved using DBCO click chemistry reaction. Enriched proteins were next subjected to DIA-MS analysis. In addition to the BONCAT enriched samples (BONCAT-ANL and BONCAT-Met), DIA-MS analysis was performed on the tumor bulk input lysates that were not subjected to BONCAT enrichment (Bulk-ANL and Bulk-Met). The proteomes of the four BONCAT-ANL samples were highly correlated among each other (Pearson’s correlation coefficient, r=0.94-0.97) (**Supplementary Fig 3A**). Principal component analysis (PCA) categorized the samples into two distinct groups; BONCAT-ANL and BONCAT-Met samples, pointing to the specificity and efficacy of ANL labeling and BONCAT enrichment (**Fig 3C**). Interestingly, the proteomes of all bulk tumor samples (four Bulk-ANL and four Bulk-Met) were highly correlated with each other (Pearson’s correlation coefficient, r=0.98-0.99) (**Supplementary Fig 3B**), and PCA was not able to differentiate Bulk-ANL from Bulk-Met samples (**Fig 3D**), suggesting that the ANL labeling does not affect the bulk tumor proteome.

Comparison of DIA-MS results from the BONCAT-ANL and BONCAT-Met samples confirmed that the majority of the proteins detected belong to the BONCAT-ANL samples, with some non-specific background present in the BONCAT-Met samples (**Fig 4A**). Among the highly enriched candidates, many proteins critical for pancreatic tumorigenesis, including KRAS, YAP1, HMGB1, HMGB2, and LEG3 (galectin-3) were identified (**Fig 4A**). There were 4405 proteins identified in the BONCAT-ANL samples (**Supplementary Table 1**). Only proteins enriched by at least two-fold in BONCAT-ANL compared to BONCAT-Met samples at a false discovery rate (FDR) < 0.05, were considered as true proteins expressed in tumor cells. Subtracting the non-specific background identified in the BONCAT-Met samples resulted in a total of 3727 BONCAT-ANL-specific proteins (**Fig 4C**). These proteins together represent the tumor cell proteome of the murine pancreatic cancer.

**Figure 4.**
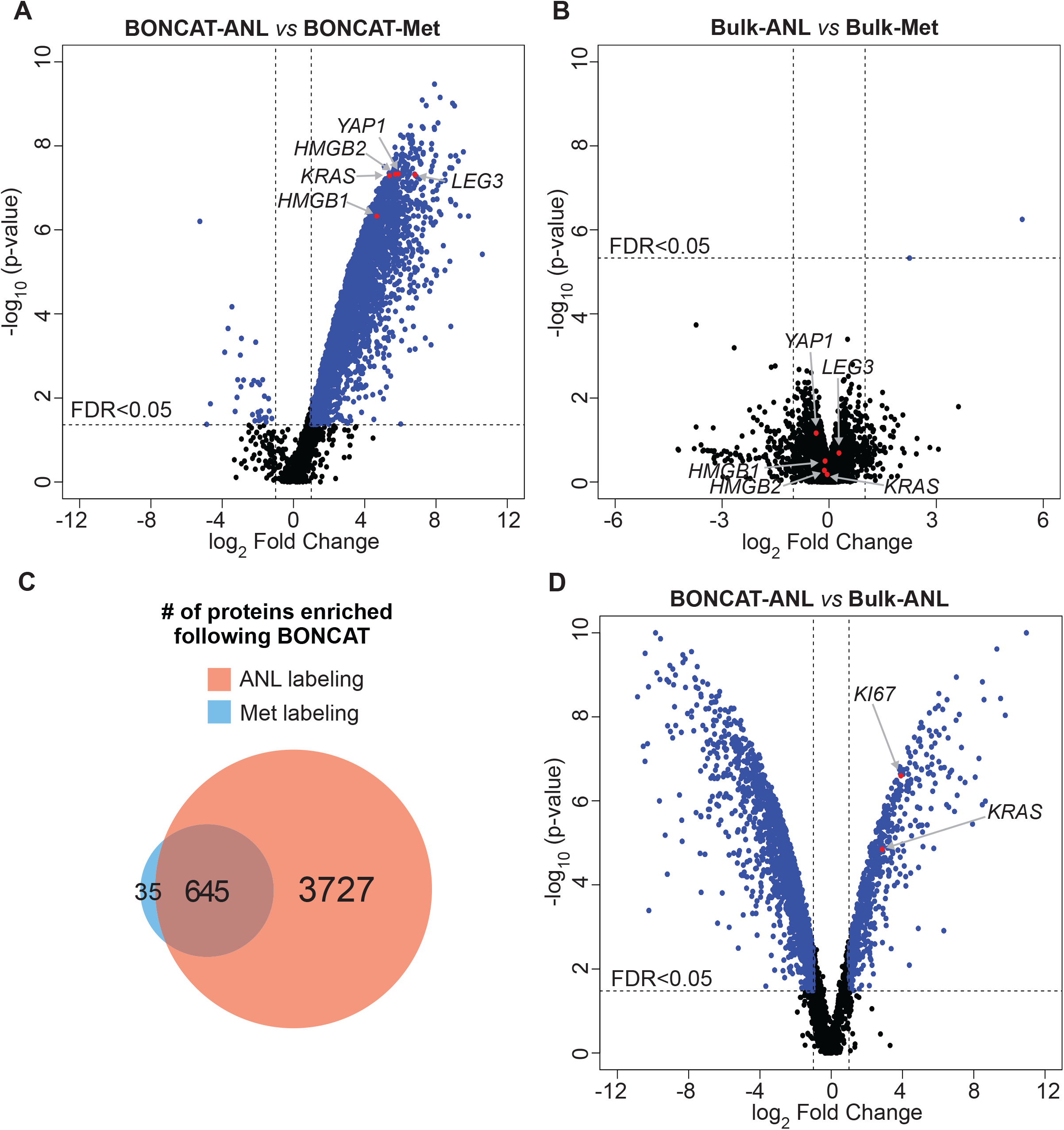
Tumor cell and tumor bulk proteomes. (A) Comparison of the BONCAT-ANL and BONCAT-Met samples. Selected proteins critical for pancreatic tumorigenesis are labeled. (B) Comparison of the bulk proteomes from ANL and Met total tumor lysates. (C) Venn diagram of the number of proteins identified by BONCAT enrichment in the ANL-labeled samples compared to the Met-labeled samples. (D) Comparison of BONCAT-ANL proteome and the corresponding Bulk-ANL proteome differentiates tumor cell specific proteins from those of non-tumor cells within the TME. Tumor specific marker proteins, such as KRAS and KI67, are highlighted in the plot.

DIA-MS proteomic analysis of the bulk tumor, comprising tumor cells, non-tumor cell types, and the extracellular matrix components, identified >5800 proteins from ANL and Met groups (Bulk-ANL and Bulk-Met), with negligible differences between the two (**Fig 4B**, **Supplementary Table 2**), confirming that the ANL labeling process does not interfere with the general protein sysnthesis machinery in PDAC tumors. Notably, the protein levels of the candidates critical for pancreatic tumorigenesis, such as KRAS, YAP1, HMGB1, HMGB2, and LEG3 (galectin-3), were not different between Bulk-ANL and Bulk-Met samples, further suggesting that the tumors from ANL and Met groups are biologically identical (**Fig 4B**). Thus, coupling BONCAT and DIA-MS allows for the exclusive *in vivo* dissection of the PDAC tumor cell proteome in a physiological context.

### Delineating proteins expressed in tumor cells from non-tumor cells within the PDAC tumor microenvironment

BONCAT-enriched proteins represent the tumor cell proteome, while the tumor bulk proteome (Bulk-ANL and Bulk-Met) encompasses the entire tumor and non-tumor proteins within the TME. Notably, many proteins expressed by the tumor cells, such as housekeeping proteins, may also be abundantly produced by other cell types within the tumor bulk. Identifying the proteins preferentially expressed in either tumor cells or non-tumor cells within the TME is critical for the study of tumor cell intrinsic carcinogenesis, dynamic interaction between tumor cells and their environment, and the discovery of novel therapeutic targets and biomarkers.

To identify tumor cell-specific and TME-specific proteins, we compared the BONCAT-enriched tumor cell proteome to the pre-enrichment tumor bulk proteome within the ANL labeled group (BONCAT-ANL *vs* Bulk-ANL). For each protein, the ratio of abundance in BONCAT-enriched to tumor bulk indicates the preferential distribution in tumor or non-tumor cells. A high BONCAT-ANL/Bulk-ANL ratio points to tumor cell-specific expression, while a low ratio implies preferential expression in various non-tumor cells within the TME (**Supplementary Table 3**). Notably, the PDAC driver oncogene KRAS (RASK) was detected as a tumor cell-specific candidate with a fold change ratio of 7.3. Another tumor cell-specific protein, KI67, was also highly enriched with a ratio of 15.1. These data provide evidence that our analysis indeed identifies tumor cell specific protein expression (**Fig 4D**). To further validate our analysis, we examined signature proteins expressed exclusively in non-tumor cells within the TME. The PDAC TME of the NSG host mice contains various cell types, such as stromal fibroblasts, monocytes/macrophages, dendritic cells, neutrophils, and endothelial cells. We, therefore, searched for the signature proteins of these non-tumor cell types in our BONCAT-ANL and Bulk-ANL data sets.^25,26^ Among 34 signature proteins present in the bulk tumor lysates, the majority of them were either not detected or at very low levels in the BONCAT-ANL tumor cell proteome. In contrast, housekeeping proteins, such as GAPDH and β-actin, did not differ between the BONCAT-enriched tumor cell proteome and the bulk proteome (**Supplementary Table 4**). Gene ontology (GO) term analysis of the tumor cell-enriched proteins revealed general, as well as pancreatic cancer-specific terms, including RNA metabolism, cell cycle/mitosis, apoptosis, chromatin remodeling, and RAS/MAPK signaling among the top terms (**Supplementary Table 5**). In contrast, GO analysis of the non-tumor cell proteins confirmed the overrepresentation of biological processes related to TME, such as immunity, extracellular matrix organization, WNT signaling, and antigen processing and presentation (**Supplementary Table 6**). These observations further support that our BONCAT-DIA-MS approach delineates proteins expressed in the tumor cells from non-tumor cells in the TME, compartmentalizing tumor and non-tumor proteins within the tumor bulk.

## Discussion

Coupling BONCAT bioorthogonal chemistry with DIA-MS proteomics analysis in an orthotopic pancreatic cancer model, we have developed an innovative technical framework that can specifically label, enrich, and identify the tumor cell proteome *in vivo*. The sensitivity and efficiency of this approach was validated through the identification of thousands of proteins expressed in pancreatic tumor cells within the tumor bulk. Comparative analysis of the BONCAT-enriched tumor cell proteome and the tumor bulk proteome facilitated the differentiation of proteins preferentially expressed in tumor cells from those of non-tumor cells within the TME.

Our approach has broad application in studies of tumorigenesis, cancer therapeutics, and biomarker discovery. Our platform may be applied to primary tumors isolated from human patients to systematically define their tumor cell proteomes. Patient-derived xenograft (PDX) models are increasingly utilized to investigate novel therapeutics and guide clinical cancer treatment.^27–29^ Following well-established protocols, primary PDX tumors can express the MetRS^L274G^ enzyme via lentiviral infection for tumor cell-specific proteomic labeling and characterization.^30,31^ Our approach, therefore, enables *in vivo* tumor cell-specific proteomic characterization in PDX models, providing an unprecedented ability for systemic interrogation of therapeutic responses at the level of individual proteins. Additionally, this technical framework may be implemented to reveal the tumor cell-specific secretome.^32^ ANL labeling of the tumor cell proteome in PDX models allows for selective purification and enrichment via BONCAT of various proteins secreted by tumor cells into the systemic circulation. Subsequent identification of the tumor cell-specific secretome using mass spectrometry will open new opportunities for the development of novel biomarkers for early cancer detection, a particularly persistent challenge in pancreatic cancers.

## Materials and Methods

### Cell lines, constructs, and chemical reagents

The 4292 murine PDAC cell line was a generous gift from Dr. Marina Pasca di Magliano.^24^ KRAS expression in this cell line is controlled by the Tet-ON system. The MetRS^L274G^ coding sequence was PCR amplified using pMarsL274G construct (Addgene 63177) as a template and inserted into BamHI and MluI sites within the pLV-EF1a-IRES-puro vector (Addgene 85132) to produce the Lentiviral-MetRS^L274G^ vector. The FLAG M2 antibody used for the detection of MetRS^L274G^ protein was purchase from Sigma. ANL, H-L-Lys(N3)-OH*HCL (HAA1625), was obtained from Peptide Solutions (Tucson, AZ). DBCO agarose beads (1034), and DBCO-TAMRA (A131), were purchased from Click Chemistry Tools and Tris [(1-benzyl-1H-1, 2, 3-triazol-4-yl)methyl] amine from Fisher Scientific.

### Click Chemistry Reactions

#### TAMRA reaction

To examine labeling efficiency, 20 μg protein lysate was incubated with 30 μM DBCO-TAMRA (absorbance/emission of 548/562 nm)/PBS (pH 7.4) for 1 hr at room temperature. Samples were boiled in Laemmli buffer and run on SDS-PAGE gel. Electrophoresed samples were visualized using the ChemiDoc imaging system (Bio-Rad) with Pro-Q Diamond filters. The gel was subsequently stained with Imperial stain (Coomassie dye R-250, Thermo Fisher 24615) according to the manufacturer’s recommendation, and visualized on the ChemiDoc imaging system.

#### CuAAC

Copper-assisted click reaction was performed on paraffin-embedded slides. Slides were deparaffinized in two changes of xylene, 5 min per change, and rehydrated sequentially in two changes, 5 min each of 100% ethanol and 95% ethanol, and 5 min 70% ethanol, and changed into water. To quench endogenous peroxidase, slides were immersed in 3% H_2_O_2_ for 15 min at room temperature. Slides were washed three times with PBS/0.1% triton X-100. The copper-assisted reaction was essentially performed as described.^33^ Briefly, the orthogonal tagging reaction was assembled in the dark; 5 μl of 200 mM TBTA, 5 μl of 500 mM TCEP, 5 μl of 2 mM biotin-alkyne-tag (Click Chemistry Tools, 1266), and 5 μl of 200 mM CuSO_4_ were added in the specified order to 5 ml of PBS (pH 7.8), and the mixture was vortexed for 10 sec after each addition. The slides were reacted with the mixture overnight at room temperature. Slides were subsequently washed three times, 20 min each, in PBS (pH 7.8), 0.5 mM EDTA, 1% Tween 20, followed by two washes, 10 min each, of PBS (pH7.8), 0.1% Tween 20. Slides were finally washed twice with PBS (pH 7.4). For signal amplification and HRP conjugation, samples were incubated with VECTASTAIN Elite ABC reagent for 30 min, washed for 15 min in PBS (pH 7.4), and changed into water. Signals were detected using ImmPACT DAB Peroxidase (HRP) Substrate (Vector laboratories, SK-4105).

#### IHC

Slides were incubated in three washes of xylene, 100% ethanol, and 95% ethanol for 5 min each. Sections were then washed in water twice, 5 min per wash. Antigens were unmasked by boiling the slides for 15 min in antigen unmasking citrate buffer (Cell Signaling Technology 14746), and cooled at room temperature. Staining was performed with VECTASTAIN ABC Elite kit according to the accompanying protocol and detected as mentioned. α-SMA antibody (19245) was purchased from Cell Signaling Technology.

### Animal models, orthotopic transplantation, and ANL labeling

Animal studies and experimental protocols were approved by the Institutional Animal Care and Use Committee at Houston Methodist Research Institute. All experimental methods were performed in accordance with the relevant national and institutional guidelines and regulations. Six-eight week old *NOD-scid IL2Rγ*^*null*^ (NSG) mice underwent surgical orthotopic injection of the pancreatic cancer cells into the pancreas. Carprofen medicated gel (5 mg/kg/day) was used for analgesia prior to the surgery, and within three days following the surgical procedure. Mice were anesthetized with isoflurane. The abdominal skin directly above the spleen was incised, the pancreas was retracted laterally and positioned outside the body. Direct injection of 1×10^6^ cells was performed using a 28.5-G needle. The needle was inserted through the knot into the pancreas tail and passed into the pancreas head to deliver the cells. Following cell injection, the spleen and pancreas were returned to the peritoneal cavity and the abdominal muscle and the skin layers were sequentially sutured. One day following surgery doxycycline was administered through the drinking water at a concentration of 0.2 g/l in a solution of 5% sucrose, and replaced every three four days. On day 4 post-surgery, experimental and control mice were IP injected with 0.1 mg/g per day of the amino acid analog or normal saline respectively, for 10 days.^34^

### BONCAT enrichment

Tumor nodules were harvested and snap frozen in liquid nitrogen until further use. Frozen tumor samples were homogenized for 20 – 60 sec in PBS (pH 7.4), 1% SDS, 100 mM chloroacetamide, and protease inhibitors. The homogenate was left at room temperature for 20 – 30 min to allow protein solubilization. Lysates were boiled for 10 min and centrifuged at room temperature at 16,000 g for 10 min. The supernatant was separated and aliquoted. Protein concentration was determined using BCA protein assay. The supernatants were used for the identification of tumor bulk (Bulk-ANL and Bulk Met) and tumor cell proteomes following BONCAT enrichment (BONCAT-ANL and BONCAT-Met). The conditions for BONCAT enrichment were based on a previously published protocol.^35^ Approximately 1.5 mg protein was diluted 2X with 8 M urea/0.15 M NaCl/PBS (pH 7.4) to a total volume of 1 ml. Fifty microliters of DBCO-agarose bead 2X slurry was washed three times with 0.8% SDS in PBS (pH 7.4). The diluted protein sample was added to the washed resin and shaken at 1200 rpm at room temperature for more than 12 hr. Unreacted DBCO was quenched by the addition of 2 mM ANL for 30 min. Resins were washed with 1 ml water and reduced with 1 mM DTT (in 0.8% SDS in PBS) for 15 min at 70°C with shaking at 1200 rpm. Free thiols were subsequently blocked with 40 mM iodoacetamide (in 0.8% SDS in PBS) for 30 min in the dark with shaking at 1200 rpm. Resins were then subjected to the following washes: 40 ml 0.8% SDS in PBS, 40 ml 8 M urea, and 40 ml 20% acetonitrile. Beads were then washed with 10% acetonitrile in 50 mM ammonium bicarbonate. The beads were spun at 2000 g to remove the liquid and resuspended in 100 μl 10% acetonitrile in 50 mM ammonium bicarbonate and 100 ng trypsin (Thermo Scientific Pierce, 90057). Beads were digested at 37°C on a shaking platform overnight, washed three times with 20% acetonitrile, and subsequently removed using centrifuge columns (Thermo Scientific, 89868). Digested peptides were dried at 30°C with vacuum concentrator (Vacufuge Plus, Eppendorf), and subsequently subjected to DIA-MS analysis (see below).

### Lysis and digestion of tumor bulk

Tumor bulk cells were lysed in a buffer containing 5% SDS/50 mM triethylammonium bicarbonate (TEAB) in the presence of protease and phosphatase inhibitors (Halt; Thermo Scientific) and nuclease (Pierce™ Universal Nuclease for Cell Lysis; Thermo Scientific). Aliquots corresponding to 100 μg protein (EZQ™ Protein Quantitation Kit; Thermo Scientific) were reduced with tris (2-carboxyethyl) phosphine hydrochloride (TCEP), alkylated in the dark with iodoacetamide and applied to S-Traps (mini; Protifi) for tryptic digestion (sequencing grade; Promega) in 50 mM TEAB. Peptides were eluted from the S-Traps with 0.2% formic acid in 50% aqueous acetonitrile, quantified using Pierce™ Quantitative Fluorometric Peptide Assay (Thermo Scientific) and diluted as needed to achieve a concentration of 0.4 μg/μl.

### DIA-MS proteomic analyses

Experimental samples were randomized for sample preparation and analysis. DIA-MS analyses were conducted on an Orbitrap Fusion Lumos mass spectrometer (Thermo Scientific). On-line HPLC separation was accomplished with an RSLC NANO HPLC system (Thermo Scientific/Dionex): column, PicoFrit™ (New Objective; 75 μm i.d.) packed to 15 cm with C18 adsorbent (Vydac; 218MS 5 μm, 300 Å); mobile phase A, 0.5% acetic acid (HAc)/0.005% trifluoroacetic acid (TFA) in water; mobile phase B, 90% acetonitrile/0.5% HAc/0.005% TFA/9.5% water; gradient 3 to 42% B in 120 min; flow rate, 0.4 μl/min. Separate pools were made of all of the samples in each experiment (equal volumes from the BONCAT-ANL/BONCAT-Met digests; equal quantities for the tumor bulk lysate digests). For the tumor bulk lysates, injections of 2 μg peptides of the pooled samples were used for chromatogram library generation.^36^ For the BONCAT-ANL and BONCAT-Met samples, aliquots of the pool of equal volumes of the digests were injected. To create the DIA chromatogram library for each sample type, the indicated peptide quantities were analyzed using gas-phase fractionation and 4-m/z windows (staggered; 30-k resolution for precursor and product ion scans, all in the orbitrap) and the MS files processed in Scaffold DIA (v2.1.0; Proteome Software) and searched against a predicted spectral library generated from the UniProt_mouse (2019_01) protein database by Prosit.^37^ Injections of 2 μg of peptides were employed for DIA-MS analysis of the individual bulk tumor lysate digests while injections corresponding to equal volumes were used for the BONCAT-ANL and BONCAT-Met samples. MS data for all individual digests were acquired in the orbitrap using 12-m/z windows (staggered; 30-k resolution for precursor and product ion scans) and searched against the chromatogram library. Scaffold DIA (v2.1.0; Proteome Software) was used for processing the DIA data from the experimental samples. Only peptides that were exclusively assigned to a protein were used for relative quantification, with two minimum peptides required for each protein and a protein-level FDR of 1%.

Correlations among different BONCAT-enriched samples and total lysate samples were analyzed by Pearson correlation. Protein intensity values (Supplementary Table 1, Supplementary Table 2) were log10-transformed. Differentially abundant proteins were analyzed by a moderated Student’s *t*-test via the limma package.^38^ A paired design was used for the BONCAT-ANL vs. Bulk-ANL analysis to account for samples being derived from the same tumor. For log2 fold change calculations, first, missing values were imputed via the “Quantile Regression for Imputation of Left-Censored data” (QRLIC) method implemented in the Scaffold DIA software. Then, the difference in medians of the log10-transformed protein intensity values between each group was computed and converted to base 2. A two-fold change cutoff and a false discovery rate (FDR) < 0.05 threshold were used to identify differentially abundant proteins. Gene ontology term enrichment analysis was performed using enrichR.^39^

## Supplementary Figure Legends

**Supplementary Fig S1.** Full-length image of the Western blot presented in Figure 2A.

**Supplementary Fig S2.** α-SMA antibody identifies stromal fibroblasts in the tumor bulk.

**Supplementary Fig S3.** Pearson correlation of DIA-MS data from the BONCAT (S3A), and the Bulk samples (S3B). The correlation coefficients are labeled above the dot plots. The dashed triangle indicates the correlation among BONCAT-ANL samples.

## Supplementary Tables

**Supplementary Table 1.** List of proteins identified in BONCAT-enriched samples by DIA-MS.

**Supplementary Table 2.** List of proteins identified in the tumor bulk lysates by DIA-MS.

**Supplementary Table 3**. Ratio of protein abundance in BONCAT-ANL versus Bulk-ANL samples.

**Supplementary Table 4**. Levels of marker proteins from tumor and non-tumor cells within the tumor microenvironment.

**Supplementary Table 5**. GO term analysis of proteins enriched in tumor cells.

**Supplementary Table 6**. GO term analysis of proteins enriched in the tumor microenvironment.

## Author Contributions

N.G.A., S.T.W. and Y.L. designed the study and developed the approach. N.G.A. carried out all cell culture, animal studies, and click chemistry experiments. S.P. and D.M. conducted the DIA-MS analysis. D.K.S., S.T.W., L.N., N.G.A., and Y.L. analyzed the data. J.C. advised the study. N.G.A. and Y.L. wrote and S.T.W. edited the manuscript. All authors reviewed and approved the manuscript.

## Code and Data Availability

The raw DIA-MS proteomic data have been uploaded to the MassIVE repository with the dataset identifier MSV000086021, and the ProteomeXchange accession number PXD021151. All code for data analysis associated with the current submission is available upon request.

## Acknowledgments

We greatly appreciate Dr. Marina Pasca di Magliano, for generously providing us the murine pancreatic cancer cell line (4292). We would also like to thank members of David Tirrell lab and Erin Schuman lab for sharing protocols on BONCAT procedures. Mass spectrometry analyses were conducted in the Mass Spectrometry Laboratory at the University of Texas Health Science Center at San Antonio. This work was supported in part by NIH K22CA207598 (Y.L.) and NIH GM008042 (D.K.S.; a grant from the UCLA-Caltech Medical Scientist Training Program). Support from the University of Texas System Proteomics Core Network for purchase of the Lumos mass spectrometer is gratefully acknowledged.

