## Supplementary figures and images for "Selective Labeling and Identification of the Tumor Cell Proteome of Pancreatic Cancer *In Vivo*"

### Supplementary Figure 1-3

**Supplementary Fig 1**

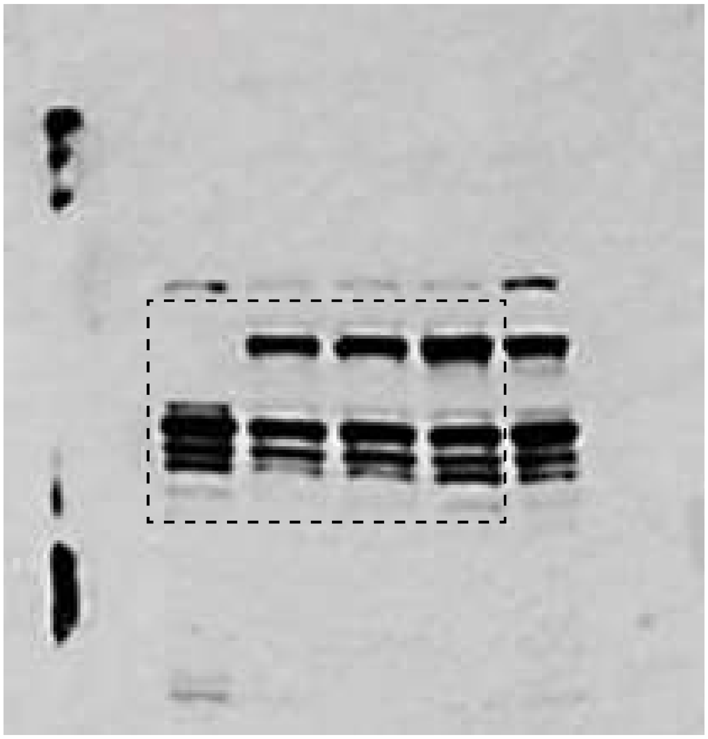

**Supplementary Fig 2**

anti-SMA (brown staining)

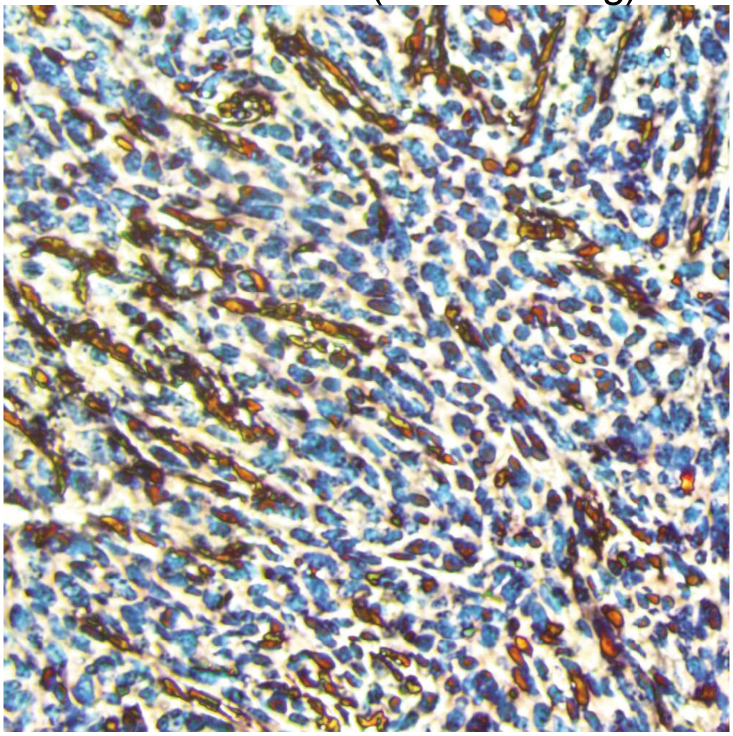

#### Supplementary Fig 3

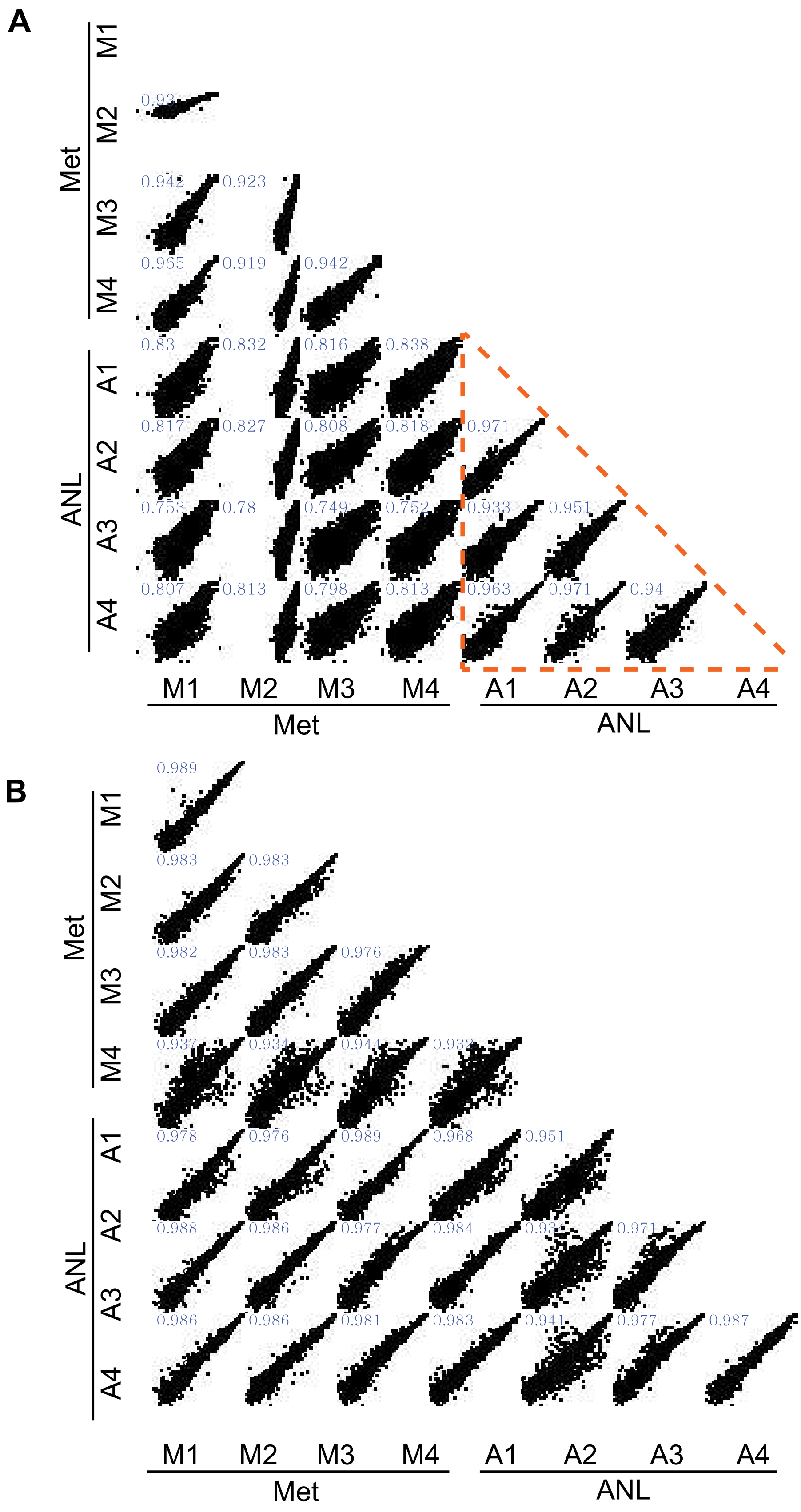
